# Effectiveness of topical antibiotics in treating corals affected by Stony Coral Tissue Loss Disease

**DOI:** 10.1101/870402

**Authors:** K.L. Neely, K.A. Macaulay, E.K. Hower, M.A. Dobler

## Abstract

Since 2014, Stony Coral Tissue Loss Disease (SCTLD) has led to mass mortality of the majority of hard coral species on the Florida Reef Tract. Following the successful treatment of SCTLD lesions on corals in closed aquaria using water dosed with antibiotics, two pastes were developed as vehicles for direct antibiotic treatments on wild corals. These pastes were tested as placebos and with additions of amoxicillin via topical applications over active SCTLD margins on multiple coral species. The effectiveness of the pastes without antibiotics (placebo treatments) was less than 10%. Adding amoxicillin to both pastes increased effectiveness. For one of the two pastes, which was silicone based with a time-release mechanism for the antibiotics, effectiveness in halting disease lesions reached 86% when amoxicillin was added. Topical antibiotic application is a viable tool for the preservation of priority corals affected by SCTLD.

## Introduction

Beginning in 2014, a disease since named Stony Coral Tissue Loss Disease (SCTLD) appeared on scleractinian corals near Miami, Florida (Precht et al. 2016). The disease is known to affect over 20 species of corals and is characterized by multifocal acute lesions which in some cases are preceded by a bleaching margin. It is highly virulent, and capable of being transmitted by physical contact as well as through seawater (Aeby et al. 2019). Progression of lesions across a colony are rapid compared to other coral diseases and in most cases, infection leads to complete mortality of the colony. Ecosystem impacts are substantial, with significant decreases in coral cover, colony density, and biodiversity recorded (Precht et al. 2016; Walton et al. 2018).

Efforts to identify the pathogen are ongoing (Meyer et al. 2019), but have not yet been successful. However, early laboratory work noted that water dosing with antibiotics resulted in disease cessation (O’Neil et al. 2018; Aeby et al. 2019). Follow up efforts by NOAA’s Coral Disease and Health Consortium (C. Woodley, pers comm) led to the development of a modified dental paste that could be applied to disease margins and is still in use by laboratories and aquariums treating SCTLD-affected corals (O’Neil et al. 2018). However, the usage of the modified dental paste requires patting the coral dry and maintaining it in low water flow for 18 hours, making it impracticable on wild corals. To resolve this, partnerships between the authors, the Florida Aquarium, and a pharmaceutical formulation and manufacturing company (Ocean Alchemists LLC and CoreRx Pharmaceuticals) led to the development of a silicone-based paste for field applications to determine whether SCTLD could be stopped on actively infected colonies at the reef.

## Methods

Corals affected with Stony Coral Tissue Loss Disease were selected for treatment at Sand Key in the lower Florida Keys. Colonies were located within a 4000 m^2^ area ranging in depth from 5 to 13 meters.

A total of 67 corals were selected for experimental treatment in October 2019. Colonies had a maximum linear dimension ranging from 12 to 432 cm. Each colony had between 1 and 12 active SCTLD lesions, and a total of 172 lesions were treated (Fig. 1). Each colony was tagged and mapped for future identification. Small masonry nails were hammered into each lesion to identify the location and progression of the disease margin.

**Figure 1.**
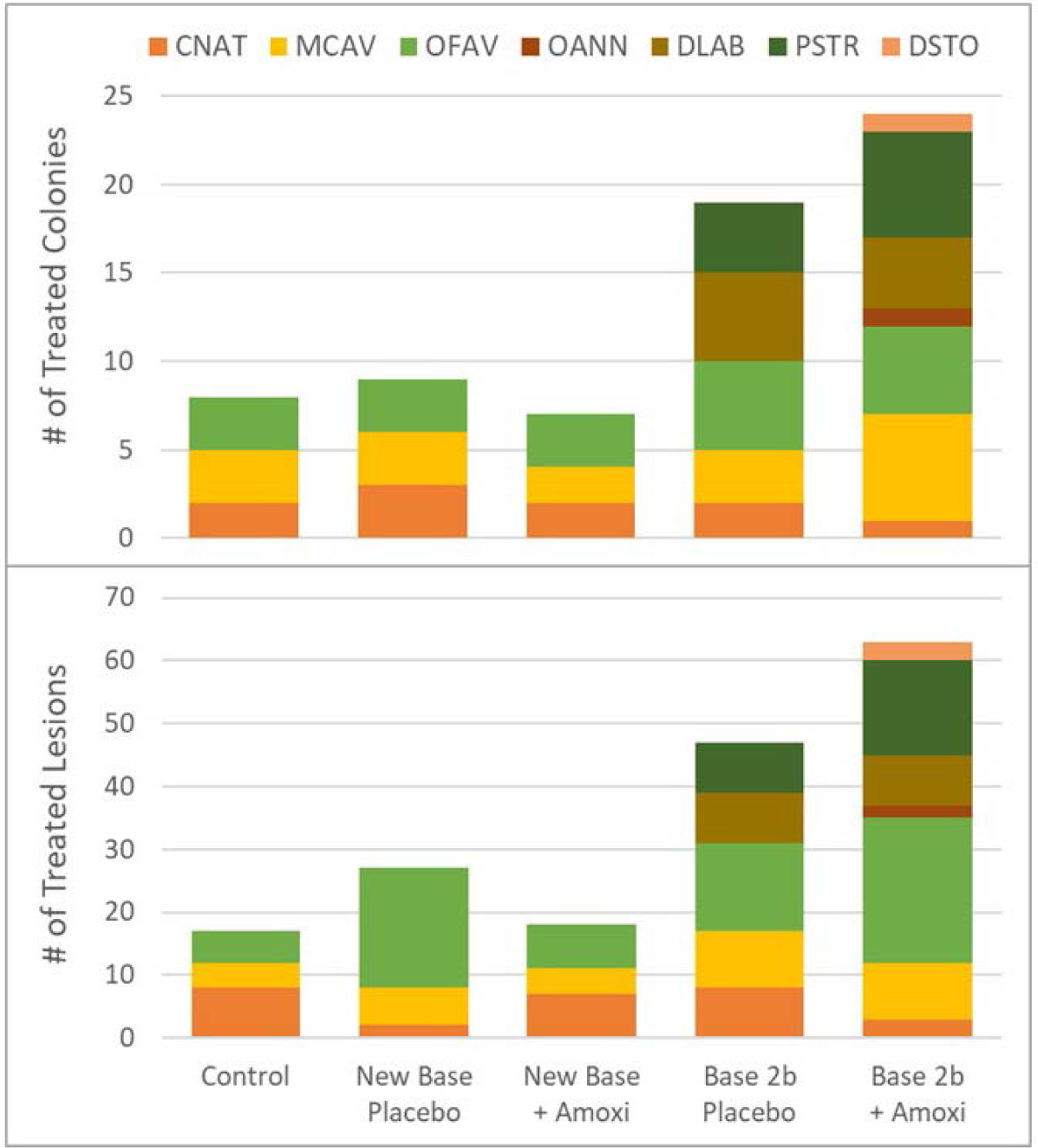
Number of colonies (top) and lesions (bottom) receiving each treatment (“Amoxi” represents addition of powdered amoxicillin). Colors show separation by species. Four-letter species codes represent: *Colpophyllia natans* (CNAT), *Montastraea cavernosa* (MCAV), *Orbicella faveolata* (OFAV), *Orbicella annularis* (OANN), *Diploria labyrinthiformes* (DLAB), *Pseudodiploria strigosa* (PSTR), *Dichocoenia stokesii* (DSTO).

Colonies were randomly assigned one of five treatments.

1. Control. Colony was tagged and nails were affixed at the disease margin, but no treatment was applied.
2. “Base 2b” Placebo. A silicone-based paste that includes polymers to mimic coral mucus consistency was applied directly to the disease margin(s).
3. Base 2b + Amoxicillin. The silicone-based paste was hand mixed with powdered amoxicillin in an 8:1 (base:amoxicillin) by weight ratio. The paste includes time-release products that regulate release of the amoxicillin over a three-day time period.
4. “New Base” Placebo. A biodegradable hydrophobic ointment designed to hold and release antibacterial compounds.
5. New Base + Amoxicillin. The New Base Placebo was mixed with powdered amoxicillin in an 8:1 by weight ratio. Release modifiers in the base facilitate amoxicillin release over three-days.

Treatments were prepared less than 6 hours before application by mixing amoxicillin into treatments 3 and 5, and by packing treatments 2-5 into 60cc catheter syringes. At each affected coral, a treatment was squeezed from the syringe and pressed onto the length of the disease margin in a band approximately 1 cm wide; approximately 0.5 cm of this anchored onto the dead skeleton while approximately 0.5 cm covered live tissue. If there were multiple lesions on a coral, they all received the same treatment.

Corals were revisited for monitoring one week and four weeks after initial treatment. At each coral, the number of effective and ineffective treatments were tallied. Photographs were also taken and arranged so before and after photos of each lesion could be compared. All analyses are based on the photographic comparisons because more lesions could be positively identified. Effectiveness was defined as the cessation of disease progression at the treatment line. Ineffectiveness was defined as the disease continuing unimpeded across the colony. Photographs were also reviewed to identify whether new lesions had developed on each colony during the post-treatment interval. After the one-month monitoring, all lesions on all surviving corals were treated with Base 2b + Amoxicillin.

## Results/Discussion

The proportion of effective treatments varied by treatment type (Fig. 2). For the controls (colonies on which no treatment was applied), 0/17 treatments were effective. Both placebo base treatments were also largely ineffective, halting progression on only 1/27 (4%) lesions treated with New Base Placebo treatment, and 4/47 (9%) lesions treated with Base 2b Placebo. The four effective Base 2b Placebo treatments consisted of three effective treatments on a single *C. natans* colony and one effective treatment on a single *D. labyrinthiformes* colony.

**Figure 2.**
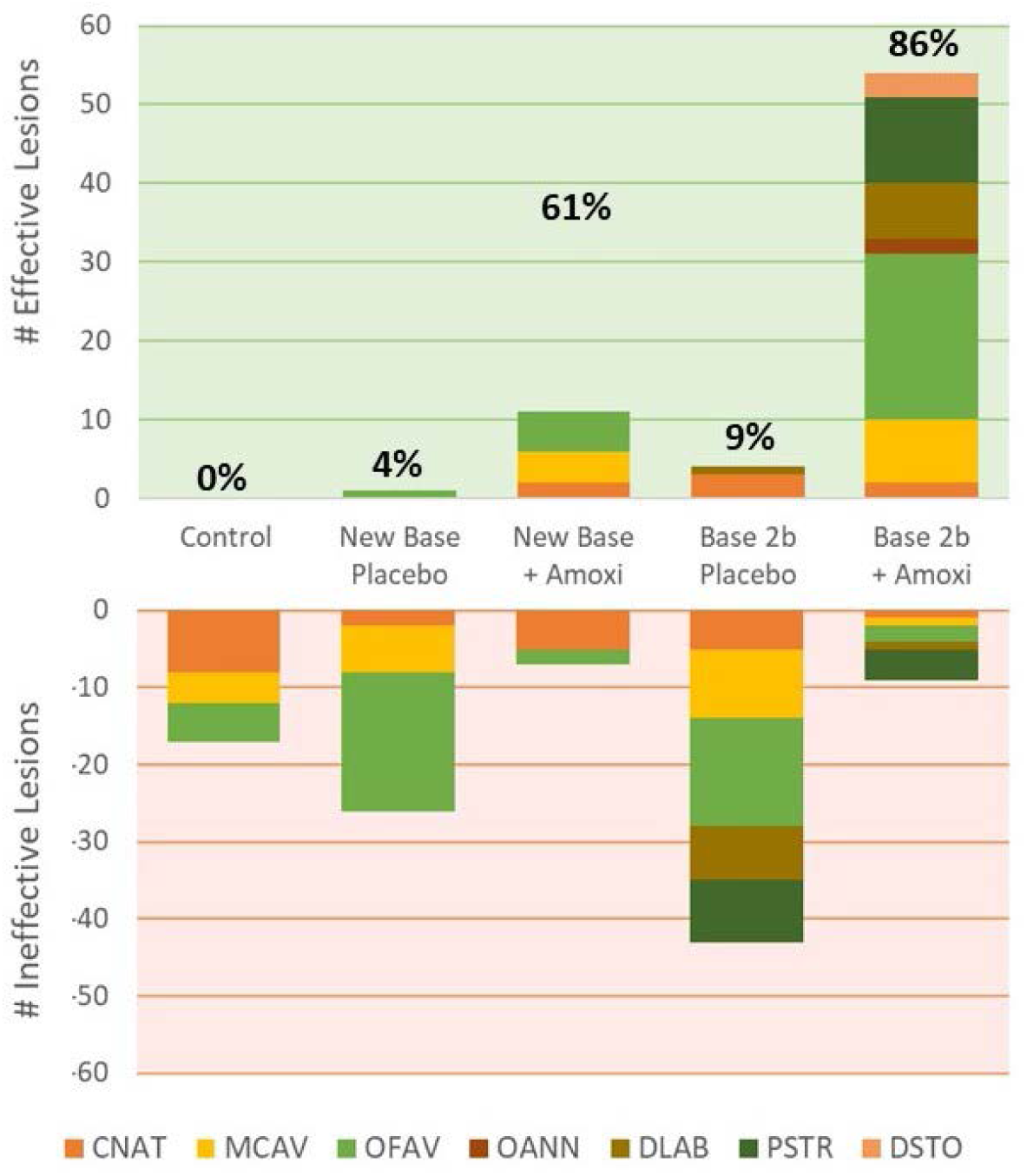
Number of effective treated lesions (top) and ineffective treated lesions (bottom) within each treatment. Colors represent different species. Total percentage of effectively treated lesions for each treatment type are shown above the effective lesion bars.

Adding amoxicillin to both placebo bases increased effectiveness. New Base + Amoxicillin effectiveness was 61% (11/18). Of the 7 failures, 5 occurred on *Colpophyllia natans*; for this species, effectiveness was only 28% (2/7). Base 2b + Amoxicillin effectiveness was 86% (54/63). Of the 9 failures, 4 occurred on *Pseudodiploria strigosa*; for this species, effectiveness was 73% (11/15).

One month after treatment, one of the control colonies and six of the Base 2b Placebo colonies had died completely. Amoxicillin-treated corals developed new lesions on 4/6 colonies treated with New Base + Amoxicillin and on 11/23 colonies treated with Base 2b + Amoxicillin.

Topical antibiotic paste was largely effective in halting disease lesions. Though laboratory antibiotic water dosing had been shown to halt SCTLD lesion progression, this trial tested and proved that a targeted and topical application was effective on wild colonies. The effect was directly tied to the addition of amoxicillin, as placebo application alone was ineffective in two different mixtures. The addition of amoxicillin to each mixture increased effectiveness by 57% (New Base) and 77% (Base 2b).

Antibiotic effectiveness is likely to remain localized within the region of application rather than spreading throughout the colony; this is evidenced by the appearance of new lesions on some amoxicillin-treated corals. To minimize mortality, coral colonies require revisitation in order to retreat any failed margins and to treat any new lesions.

Treatment of SCTLD-infected colonies using topical amoxicillin paste is an option for SCTLD disease intervention. Though requiring an investment of time and resources for initial treatment as well as repeated visitation, it is a valuable tool for preserving high-value corals.

## Acknowledgements

Funding for these activities was provided by Florida’s Department of Environmental Protection. Work was conducted in the Florida Keys National Marine Sanctuary under permit FKNMS-2019-115.

## Literature Cited

Aeby G, Ushijima B, Campbell JE, Jones S, Williams G, Meyer JL, Hase C, Paul V (2019) Pathogenesis of a tissue loss disease affecting multiple species of corals along the Florida Reef Tract. Frontiers in Marine Science 6:678

Meyer JL, Castellanos-Gell J, Aeby GS, Häse CC, Ushijima B, Paul VJ (2019) Microbial Community Shifts Associated With the Ongoing Stony Coral Tissue Loss Disease Outbreak on the Florida Reef Tract. Frontiers in Microbiology 10

O’Neil K, Neely KL, Patterson J (2018) Nursery management and treatment of idsease-ravaged pillar coral (*Dendrogyra cylindrus*) on the Florida Reef Tract. Florida DEP, Miami, FL 13

Precht WF, Gintert BE, Robbart ML, Fura R, van Woesik R (2016) Unprecedented Disease-Related Coral Mortality in Southeastern Florida. Scientific Reports 6

Walton CJ, Hayes NK, Gilliam DS (2018) Impacts of a Regional, Multi-Year, Multi-Species Coral Disease Outbreak in Southeast Florida. Frontiers in Marine Science 5

